# Stepwise molecular specification of excitatory synapse diversity on a target neuron

**DOI:** 10.1101/2023.01.03.521946

**Authors:** Maëla A. Paul, Séverine M. Sigoillot, Léa Marti, Marine Delagrange, Philippe Mailly, Fekrije Selimi

**Affiliations:** Center for Interdisciplinary Research in Biology (CIRB), College de France, CNRS, INSERM, Université PSL, Paris, France; Plateforme qPCR-HD-GPC, Institut de Biologie de l’Ecole Normale Supérieure, Paris, France

**Author notes:** Equal contribution (Maëla A. Paul, Séverine M. Sigoillot).

**Keywords:** Brain, development, synaptogenesis, excitatory synapse, cerebellum, Purkinje cell, inferior olivary neuron, granule cell, transcriptomics, mouse

## Abstract

Brain function relies on the generation of a large variety of morphologically and functionally diverse, but specific, neuronal synapses. Here, we show that, initially, synapse formation on a common target neuron, the cerebellar Purkinje cells, involves a presynaptic secreted protein common for all types of excitatory inputs. The molecular program then evolves only in one of the inputs with the additional expression of a combination of presynaptic secreted proteins that specify the mature pattern of connectivity on the target. These results show that some inputs actively and gradually specify their synaptic molecular identity while others rely on the “original code”. Thus, the molecular specification of excitatory synapses, crucial for proper circuit function, is acquired in a stepwise manner during mouse postnatal development and obeys input-specific rules.

## Introduction

Morphologically, molecularly and functionally distinct types of neurons need to be connected in a stereotyped manner to form the short-range and long-range circuits that underlie complex behaviors in mammals. This requires not only the recognition of the proper neuronal partners, *via* the molecular control of cell migration and axon guidance in particular, but also the formation of specific synapses on a given target neuron^1,2^. Each neuron receives multiple types of inputs that form synapses with specific identities determined by their subcellular localization and their degree of connectivity on the target neuron, as well as their specific molecular composition and functional properties. Indeed, synaptome mapping using two excitatory postsynaptic markers and unsupervised classification has revealed thirty seven synaptic types^3^. Given that more than a thousand synaptic proteins have been identified, the mammalian brain thus likely contains a large diversity of synapses^3^ supporting information processing, learning and memory.

Various principles have been proposed to control the specific wiring of neurons during development^1,2,4–6^. Neuronal activity sculpts circuits *via* selective stabilization or elimination of synapses^7^. On the other hand, the chemoaffinity hypothesis states that the specificity of connectivity is regulated by “specific cytochemical affinities”^8^ and implies the existence of different molecular combinations for each synapse type. While the chemoaffinity hypothesis has received much attention in the case of neuronal partner selection, to what extent it also relates to synapse specificity on a given target neuron remains to be demonstrated^2^. Many adhesion complexes of transmembrane proteins with roles in synaptogenesis have been identified (see^2,6^ for review). Different surface proteins have been shown to underlie excitatory *versus* inhibitory synapse formation^4,6,9–15^. However, synapse diversity goes well beyond these two broad functional classes and generally, a given neuron receives synapses from several types of glutamatergic or inhibitory inputs. While individual molecular complexes have been identified to participate in the formation of certain synapses^9,10,16^, whether and how a combination of proteins controls the formation of a specific synapse type remains to be tested. In addition, whether the rules underlying the specification of different excitatory synapse types on a given neuron during development are identical remain unknown.

Cerebellar Purkinje cells (PCs) are a perfect example of the type of specific and stereotyped connectivity that must be generated for proper network function and information coding. They are at the center of computation of the cerebellum^17^, an organ that is well known for its control of motor coordination and learning, but also increasingly recognized for its broader role in many cognitive processes^17,18^. PCs receive direct synapses from two excitatory inputs, the parallel fibers (PFs) and climbing fibers (CFs), axons of cerebellar granule cells (GCs) and inferior olivary neurons (IONs) of the brainstem respectively. Their activity is further modulated by local inhibitory neurons, the stellate cells and basket cells. The PF and CF synapses are made on separate subcellular territories on a given PC, distal dendritic spines versus proximal dendrites respectively, with a different degree of connectivity, and different functional properties. This mature connectivity is established during the first three postnatal weeks in rodents^19^. The formation of each excitatory synapse type depends on signaling pathways that involve a different member of the C1Q protein family^9^. Parallel fiber/Purkinje cell (PF/PC) synapses are dependent on the formation of a tripartite complex between presynaptic neurexin-1, secreted CBLN1 and postsynaptic GluD2 receptors^11,20^, while climbing fiber/Purkinje cell (CF/PC) synapses rely on secreted C1QL1 and postsynaptic adhesion GPCR BAI3^9,21^. Both CBLN1 and C1QL1 loss of function (LOF) experiments result in the loss of about half of PF/PC or CF/PC synapses, respectively^9,22^. Thus, either compensation by other synaptic proteins can prevent complete loss of synapses, or several molecular signaling pathways might be at work at each synapse type, in support of the chemoaffinity hypothesis.

Using the cerebellar PCs as a model, we searched for the combination of membrane and secreted proteins underlying the specificity of excitatory connectivity on a single neuron type. Neuron-specific gene expression profiling and LOF analysis during postnatal development revealed that the CF relies on a combination of cell surface proteins belonging to different signaling pathways for its mature connectivity on PCs. These proteins, in particular LGI2 and CRTAC1, could play a function similar to the one of C1QL1, the well-known CF/PC synaptogenic molecule. Surprisingly, we discovered that the cerebellar GC marker *Cbln1* is also expressed by IONs and that, in addition to its known role at PF/PC synapse, it also regulates synaptogenesis between CFs and PCs at early stages. Thus, a common molecular code is shared initially, and a specific molecular combination is acquired progressively in selected input types during postnatal development to establish the final connectivity pattern on a given target neuron.

## Results

### Purkinje cell excitatory inputs are characterized by different surfaceome expression dynamics during postnatal development

Synapse specificity and identity involves the recognition and adhesion between the pre- and post-synaptic domains of neuronal partners. Thus, molecules that regulate synapse formation and specification are expected to be part of the “surfaceome”, defined as the set of genes coding for membrane and secreted proteins. In a previous study^23^, neuron-specific expression profiling *in vivo* and comparative analysis identified the differentially expressed genes (DEGs) specific for each adult PC excitatory input (401 for cerebellar GCs versus 598 for IONs of the brainstem). Amongst these DEGs, 74 and 250 genes code for membrane and secreted proteins and thus constitute the specific adult surfaceome of GCs and IONs, respectively (Figure 1A). As expected, these genesets contained genes coding for proteins with known function in the formation and stabilization of excitatory synapses on PCs: *Cbln1* for synapses from GCs and *C1ql1* for synapses from IONs. Because the pattern of expression of *Cbln1* and *C1ql1* is highly correlated with the respective timing of synaptogenesis on PCs (Figure 1A), we reasoned that other genes with similar roles in synapse specification should have similar expression pattern dynamics during development. High-throughput expression analysis of the GC and ION genesets was performed at different postnatal developmental stages using RNA extracts from the cerebellum and brainstem (supplemental data 1 and 2): from E17, a time when IONs and PCs have already been generated and have started to connect, to adult when all the connectivity between GCs and IONs onto PCs has been established^19^. Clustering analysis showed a fundamental difference in the postnatal dynamics of the surfaceome that characterizes each PC excitatory input (Figure 1B and 1D). In GCs, only one major cluster was identified containing 70 genes with a developmental pattern similar to the one of the synaptogenic gene *Cbln1*: they were expressed at very low levels during the first two postnatal weeks, and then, expression synchronously increased starting at P14 (Figure1B), in correlation with the timing of synaptogenesis between GCs and PCs (Figure 1C). In the brainstem, clustering analysis of ION-specific surfaceome resulted in two groups: a group of 117 genes that were expressed at higher levels during the two first postnatal weeks and then decreased (cluster 1, Figure 1D), and a second group of 125 genes, which included *C1ql1*, that were expressed at increasingly higher levels starting during the second postnatal week (cluster 2, Figure 1D). Because this developmental period is essential for the establishment of mature ION-PC connectivity (Figure 1C), we searched amongst the 125 genes of cluster 2 for candidates that would play a functional role similar to *C1ql1*. Fifteen genes were selected based on high expression levels (Figure S1A) and database and literature curation looking for predicted structural domains with roles in recognition. The correlation of their expression pattern with the synaptogenic gene *C1ql1* was then determined using single molecule Fluorescent *In Situ* Hybridization (smFISH) in the inferior olive region at three ages of interest. At P4, CF, but not PF synaptogenesis, has begun on PCs with several CFs synapsing on their somata. At P14 the single wining CF has already started to translocate^24^ and CF and PF inputs compete for their subcellular synaptic territory on the growing PC dendrites^25^. Finally, the adult mature stage is reached when the final territory of innervation and functional properties of excitatory synapses on PCs have been acquired. Interestingly, computation of a pixel-based Pearson correlation coefficient (see methods section) allowed the classification of the candidate genes in groups that corresponded to the different developmental stages of ION-PC connectivity. Four genes, *Nrcam, Lgi2, Crtac1* and *Sema4f*, were highly correlated with *C1ql1* (Pearson coefficient > 0.6) at every stage (Figure 1E), suggesting a synaptogenic role similar to *C1ql1* during the postnatal development of ION-PC connectivity. Five genes (*Shisal1, Thy1, Adam11, Crh* and *Gpr123*) were less correlated at P4 but highly correlated starting at P14 (Figure 1E), suggesting a role at later stages than *C1ql1*, potentially in the functional maturation of CF/PC synapses. The other six candidates (*Tmem184b, Fstl1, Cx3cl1, Adam23, Cd151, Tmem179*) did not reach the threshold of correlation with *C1ql1* in the adult (Figure S1B), and might play roles in other processes than the control of CF/PC synaptogenesis and functional maturation. Overall, our high-throughput RTqPCR and smFISH analyses reveal a multi-step postnatal acquisition of the adult gene expression pattern specific for IONs for the combination of genes coding secreted and membrane proteins. Thus, the acquisition of the adult-specific surfaceome characterizing the two PC excitatory inputs, GCs and IONs, follows different time courses during postnatal development. GCs acquire their “surface” adult identity in one step during their differentiation, while the surfaceome of IONs changes progressively to acquire the adult combination. Indeed, a first set of genes (Figure 1D, cluster 1) is highly, but transiently, expressed during the first stage of immature ION connectivity on PC somata and the reduction of CF/PC multi-innervation by synapse elimination. The expression of a second set of genes (Figure 1D, cluster 2) increases during the second postnatal week. This set comprises genes that will participate in the control of CF translocation, synaptogenesis on PC dendrites and competition with PFs (Figure 1C), and other genes that will participate later in the functional maturation of the CF/PC synapses like *Crh*^26^ (Figure 1E, lower correlation with *C1ql1* at the single cell level at P4).

**Figure 1.**
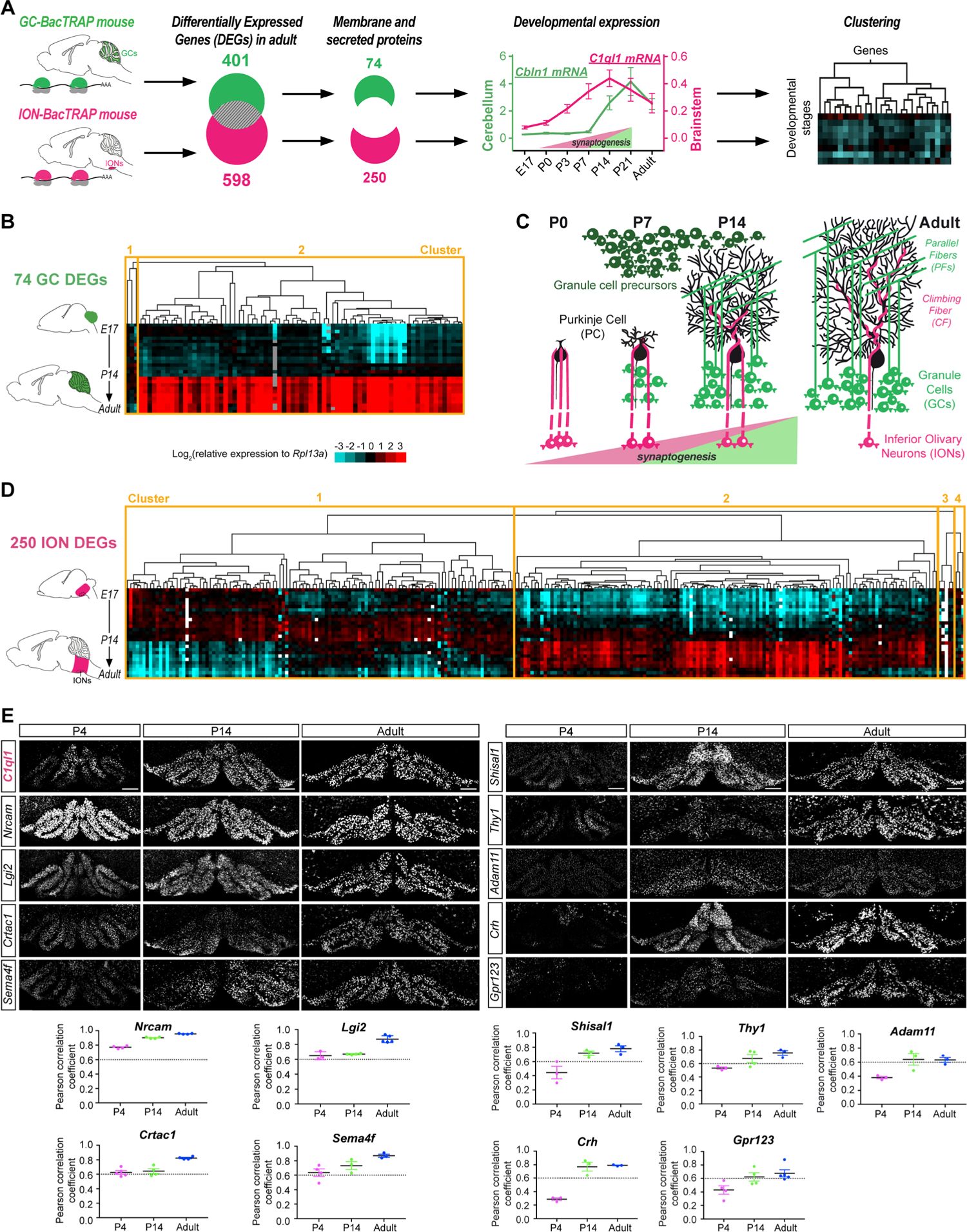
Purkinje cell excitatory inputs are characterized by different surfaceome expression dynamics during postnatal development. **(A)** Workflow for the identification of candidate synaptogenic genes for each input type. A previous analysis using neuron-specific BacTRAP mouse lines identified the adult gene expression profile specific for GCs or IONs, including 74 and 250 DEGs coding for membrane and secreted proteins, respectively^23^. The developmental expression patterns of GC and ION surfaceomes were assessed at different stages using high-throughput RT-qPCR on cerebellar and brainstem RNA extracts (Embryonic day, E17; Postnatal days, P0, P3, P7, P14 and P21; and adult). Developmental expression profiles of *Cbln1* and *C1ql1* are shown as references for synaptogenic molecules for GCs and IONs, respectively. Clustering analysis was then performed using the Spearman rank correlation method to group the genes according to their pattern across developmental stages. (**B)** *Left panel:* Heatmap of the 74 GC DEGs in the cerebellum between E17 and adult. *(C):* Schematic illustration of postnatal development and maturation of GC and ION connectivity on PC in the cerebellar cortex. **(D)** Heatmap of the 250 ION DEGs in the brainstem between E17 and adult. **(E)** Duplex smFISH for *C1ql1* and candidate mRNAs in coronal sections from brainstem at P4, P14 and in adult. The degree of correlation of expression between *C1ql1* and candidate mRNAs in the brainstem was determined by computing the Pearson correlation coefficient (coefficient >0.6 corresponding to high correlation). *Left panel:* candidates highly correlated with *C1ql1* at all developmental stages. *Right panel:* candidates highly correlated with *C1ql1* from P14 to adult. Data are presented as mean ± SEM. n = 3-5 animals per condition, 2-4 independent experiments. Scale bars, 150 µm. See also Figure S1.

### A combination of cell surface proteins controls the establishment of Climbing fiber/Purkinje cell connectivity

If synapse specification is controlled on a given target by a specific combination of presynaptic recognition proteins, the combination coding for CF/PC synapse development should include other ION DEGs with an expression pattern highly correlated to *C1ql1* at the single cell level. Using a second analysis pipeline (cf. material and methods), we thus analyzed smFISH experiments at the single-cell level (Figure 2A and B) and computed the Pearson correlation coefficient of *Nrcam, Lgi2, Crtac1* or *Sema4f* with *C1ql1* in IONs. This analysis confirmed very high correlation for *Nrcam, Lgi2* and *Crtac1* at every stage, but showed a relatively lower correlation for *Sema4f* transiently at P14 (Figure 2B). Interestingly, *Crtac1* and *Lgi2* both code, like *C1ql1*, for extracellular matrix proteins, while *Nrcam* codes for an Ig domain containing single-pass transmembrane protein (Figure 2C). While C1QL1 is characterized by a globular C1q domain known to interact with the BAI3 synaptic adhesion-GPCR^21,27^, LGI2 is part of a family of secreted proteins containing Leucine Repeat Rich domains and a beta-propeller domain (Figure 2C), and binds to the ADAM family of synaptic receptors^28^. CRTAC1 is also part of the beta-propeller domain containing family of proteins (Figure 2C), and binds to NOGO receptors^29^ that have been recently shown to promote synaptogenesis in addition to neuronal morphogenesis^30^.

**Figure 2.**
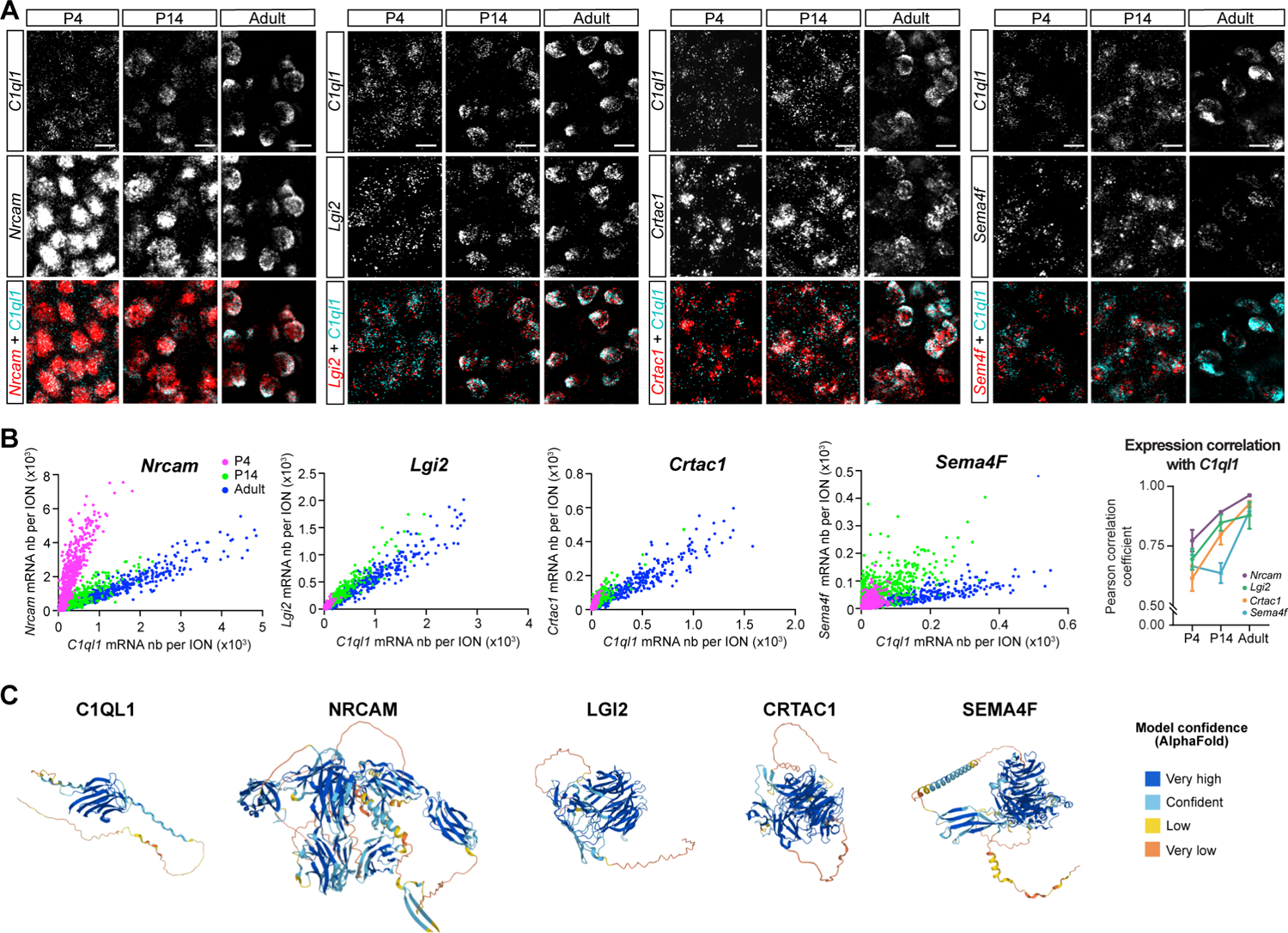
A combination of cell surface molecules characterizing IONs during postnatal development. **(A)** Duplex smFISH for *C1ql1* mRNAs and the four highly correlated candidate mRNAs, *Nrcam, Lgi2, Crtac1* and *Sema4f* in IONs at P4, P14 and in adult. **(B)** Quantification of the number (nb) of *C1ql1* and candidate gene mRNAs per ION for *Nrcam, Lgi2, Crtac1* or *Sema4f* at P4, P14 and in adult. n = 568-682 cells (1 cell corresponding to 1 dot on the graphs), one representative experiment is shown. Correlation between the expression of *C1ql1* and candidate mRNAs was determined at the single cell level (Pearson correlation coefficient >0.6 corresponding to high correlation). Data are presented as mean ± SEM. n = 3-6 animals, 2-4 independent experiments. **(C)** AlphaFold structure prediction of C1QL1 and candidate proteins NRCAM, LGI2, CRTAC1 and SEMA4F (Database: https://alphafold.ebi.ac.uk/).

To uncover the function of *Nrcam, Lgi2* and *Crtac1* in excitatory synaptogenesis and test their involvement in synapse specification on PCs, we set up an *in vivo* assay to perform ION-specific LOF of each candidate gene during postnatal development. To enable CRISPR/Cas9 based genome editing specifically in IONs during postnatal development, we developed an intersectional strategy by injecting in the cerebellum of Cas9 knock-in P0 mouse pups^31^ a retrograde AAV driving the expression of the CRE recombinase under the CamKII promoter, which is not expressed in GCs, thus enabling conditional expression of CAS9 and soluble GFP specifically in IONs (Figure 3A and B). Several guide RNAs were selected for each candidate gene using CRISPOR web-based tool, except the *Nrcam* gRNA that was previously characterized^32^ (Figure S2). Transduction of cultured cortical neurons was used to assess efficiency and specificity using sequencing and RT-qPCR (Figure S2, supplemental data 3 and material and methods). No mutation in protein coding regions was detected by sequencing, and RTqPCR showed a 50% (*Nrcam* and *Lgi2*) and 75% *(Crtac1)* decrease in mRNA levels for each of our targeted candidate. No effects on the expression of other candidate genes analyzed in our study was detected. The more efficient guide RNA was selected for each candidate gene, subcloned in the AAV construct together with the CamKII-Cre to simultaneously drive the expression of guide RNAs under the control of the U6 promoter and the conditional expression of CAS9 to edit the genomic region of interest in IONs. Consequences of ION-specific LOF of each candidate gene were assessed at P21, an age when the final pattern of connectivity between PCs and its excitatory inputs has been attained. Thanks to the co-expression of soluble GFP with CAS9 after recombination, we could readily identify CFs from genome edited IONs in the cerebellar cortex (Figure 3B). VGLUT2 is a specific marker of CF/PC synapses in the cerebellum and deficits in CF/PC synapse numbers and function have been shown consistently to be accompanied by a decrease in VGLUT2 cluster extension or numbers^9,33^. We thus used anti-GFP and anti-VGLUT2 immunolabeling followed by high resolution 3D imaging using spinning disk confocal microscopy, to quantify the morphology and the extension of the transduced GFP expressing (GFP+) CFs and, using a custom plugin, the number, and size of associated VGLUT2 presynaptic boutons (Figure 3B and C, and methods). CF branching was significantly diminished in the case of *Nrcam* LOF, but not for the other candidates and total cumulative length of the CF was unchanged (Figure 3C). In controls, CFs extend to about 70% of the molecular layer height, a value that is coherent with previously published data^9^. Small but significant changes in this extension were detected for all three candidates: *Lgi2* and *Crtac1* LOF led to a 11% and 6% decrease respectively, while *Nrcam* LOF led to a 6% increase (Figure 3C). Thus, amongst the three candidates, NRCAM plays a specific role in CF morphogenesis promoting their branching along the PC dendrites. At the synaptic level, an increase in about 17% in the number of VGLUT2 clusters per GFP labelled CF was observed after *Nrcam* LOF (mean 105.9 ± SEM 4.705 *versus* 123.7 ± 4.731, p = 0.012, Student’s unpaired t test), accompanied by a significant 9% decrease in the mean volume of those clusters (mean 1.068 ± SEM 0.038 *versus* 0.9694 ± 0.034, p = 0.029, Mann-Whitney test). On the contrary, *Lgi2* LOF and *Crtac1* LOF led to a 20% and 22% significant decrease in the mean VGLUT2 number per transduced CF, respectively (*Crtac1*: mean134.9 ± SEM 5.001 *versus* CTL: 104.8 ± 5.889, p<0.001, Student’s unpaired t test; *Lgi2*: mean 112.9 ± SEM 4.504 *versus* CTL: 90.85 ± 4.399, p<0.001, Student’s unpaired t test). This decreased mean number of presynaptic boutons was accompanied by a tendency to decreased mean volumes for *Lgi2* LOF and a 15% significant increase for *Crtac1* LOF (Figure 3C). Thus, LGI2 and CRTAC1 both promote, while NRCAM inhibits, CF/PC synaptogenesis during the second developmental phase corresponding to CF synaptogenesis on PC dendrites and competition with PFs. NRCAM plays an additional role in controlling CF morphogenesis in accordance with expression data showing a precocious strong expression of *Nrcam* in IONs (Figure 2A and^34^). LGI2 and CRTAC1 are thus synaptogenic molecules forming together with C1QL1 a combinatorial secreted code promoting CF/PC synapses.

**Figure 3.**
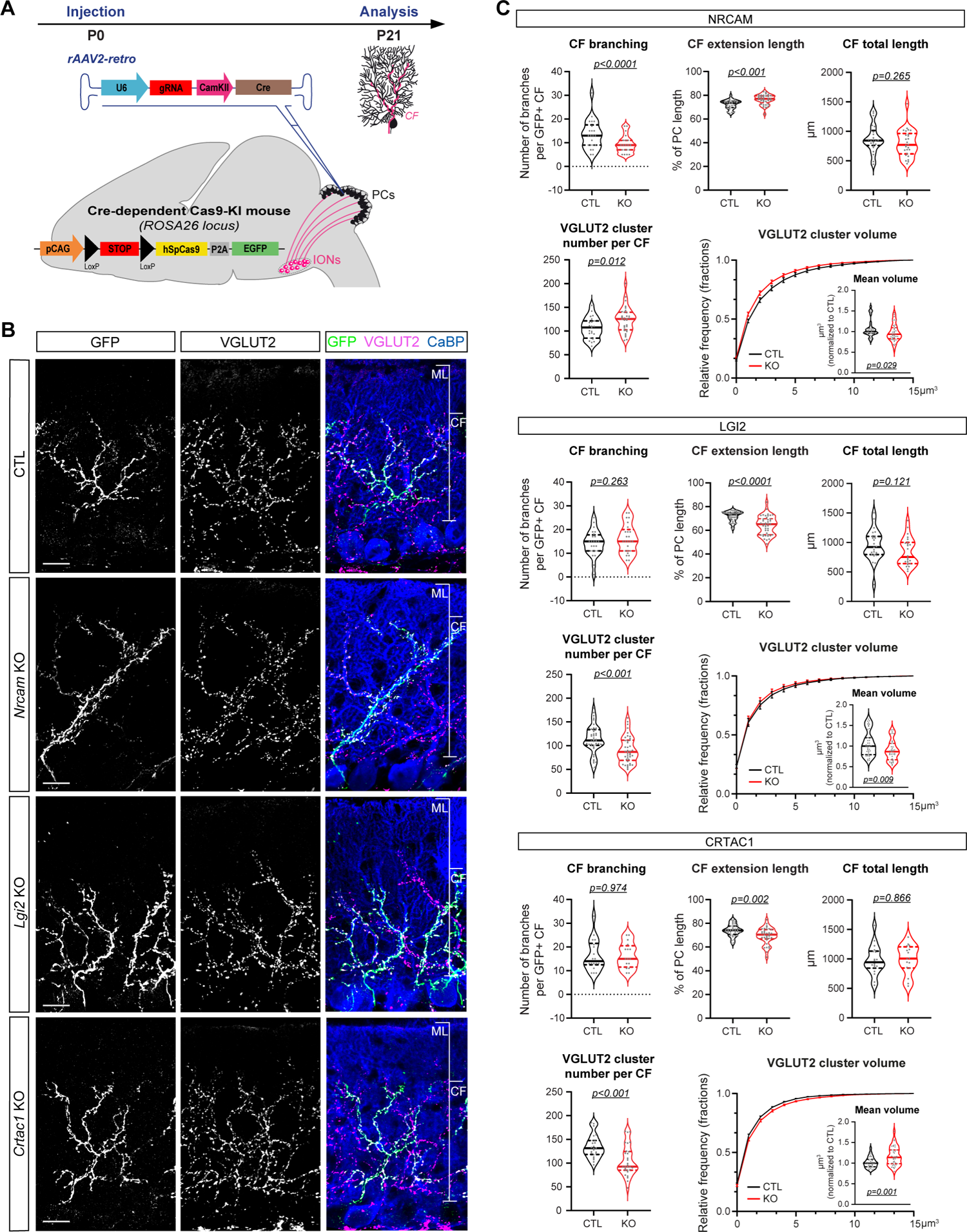
A combination of cell surface proteins controls the establishment of Climbing fiber/Purkinje cell connectivity. **(A)** Schematic illustration of the strategy of CRISPR/Cas9-mediated loss of function of candidate genes specifically in IONs. P0 stereotaxic injection of retrograde rAAVs expressing guide RNA (gRNA) targeting a candidate gene or a non-targeting control gRNA (CTL), under the U6 promoter, as well as the Cre recombinase, under the CamKII promoter, were performed in the cerebellum of Cre-dependent Cas9 knock-in (KI) mice. Analyses of the morphology and connectivity of GFP positive (GFP+) CFs (from Cas9-recombined IONs) were performed at P21 in parasagittal cerebellar sections from KO or CTL injected animals. **(B)** GFP+ CF and CF presynaptic boutons were immunostained for GFP (green) and VGLUT2 (magenta), respectively, and immunostaining for calbindin was used to label PCs and their dendritic tree (blue) in CTL and *Nrcam, Lgi2* or *Crtac1* KO. Molecular layer (ML) and CF extensions are indicated on the merge image. Scale bars, 20 µm. **(C)** GFP+ CF morphology was assessed measuring 3 parameters: the number of branches per CF, the percentage of CF extension length on PC dendrite and the total length of each CF. VGLUT2 cluster number, and distribution and mean of the VGLUT2 cluster volume were also quantified in GFP+ CFs. Data are presented as violin plot with the median and quartiles. CF morphology and VGLUT2 cluster number: n = 16-26 CFs per condition, 5-6 animals per condition; VGLUT2 cluster volume: n ≥ 33 images per condition, 9-10 animals per condition; 3-5 independent experiments. Statistics: unpaired Student’s test for CF branching (*Lgi2* or *Crtac1* KO), CF extension (*Nrcam* or *Crtac1* KO), CF total length, VGLUT2 cluster number, VGLUT2 cluster volume (*Crtac1* KO) and Mann-Whitney test for CF branching (*Nrcam* KO), CF extension (*Lgi2* KO), VGLUT2 cluster volume (*Nrcam* or *Lgi2* KO). See also Figure S2.

### Cbln1 expression in inferior olivary neurons is necessary for proper climbing fiber connectivity on Purkinje cells

Our developmental expression analysis showed a multi-step determination of ION identity with a set of cell surface molecules that are expressed highly during the first two postnatal weeks, when CF synapses are made on the somata of PCs (cluster1, Figure 1C). Surprisingly, by analyzing all 324 DEGs including those that are specific to cerebellar GCs, we found *Cbln1* in the cluster of genes highly expressed in the brainstem during the first two postnatal weeks (cluster 1, Figure S3A and supplemental data 4). *Cbln1* is a well-known marker of PF/PC synapses and is essential for their formation and stabilization^35^. We confirmed expression of *Cbln1* in IONs using smFISH: our quantitative analysis shows that *Cbln1* mRNA numbers in single IONs are comparable to the number of *C1ql1* mRNAs at P4 but then start to diminish by P14 while *C1ql1* expression on the contrary increases at the single cell level (Figure 4A). This raised the question of the role of *Cbln1* in IONs and prompted us to perform ION-specific *Cbln1* LOF at P0 and analyze the consequences on CF/PC innervation (Figure 4B). At P7, CFs lacking *Cbln1* function had started to translocate in about 29% of contacted PCs in comparison to 11% for control GFP+ CFs. *Cbln1* LOF also led to a 52% significant decrease of the mean number of VGLUT2/GFP clusters on PC somata compared to controls (mean 0.94 ± SEM 0.1038 *versus* 0.45 ± 0.0667, p = 0.029, p<0.001, Student’s unpaired t test). No effect on the mean VGLUT2 cluster volume was detected (Figure 4C and S3C). When analyzed at P14, *Cbln1* LOF led to an increase in CF branching and decreased CF extension. The total cumulative length of CF is not changed by loss of *Cbln1*. This decreased extension of the territory of innervation along the PC dendritic arbor is accompanied by a decrease in synaptogenesis since a significant 21% decrease in VGLUT2 cluster numbers on PC dendrites was also observed, with no change in the mean VGLUT2 cluster volume, after *Cbln1* LOF (Figure 4D and S3C). These results show that CBLN1 plays several roles during the establishment of CF/PC connectivity. It prevents precocious translocation of the winning CF on the growing PC dendrites, controls CF morphogenesis by directing its branching, and promotes CF synaptogenesis both on PC somata and dendrites. Given that CBLN1 is a secreted protein of the C1q family, these results finally establish CBLN1 as a common synaptogenic secreted factor for both types of excitatory inputs, GCs and IONs, connecting Purkinje cells.

**Figure 4.**
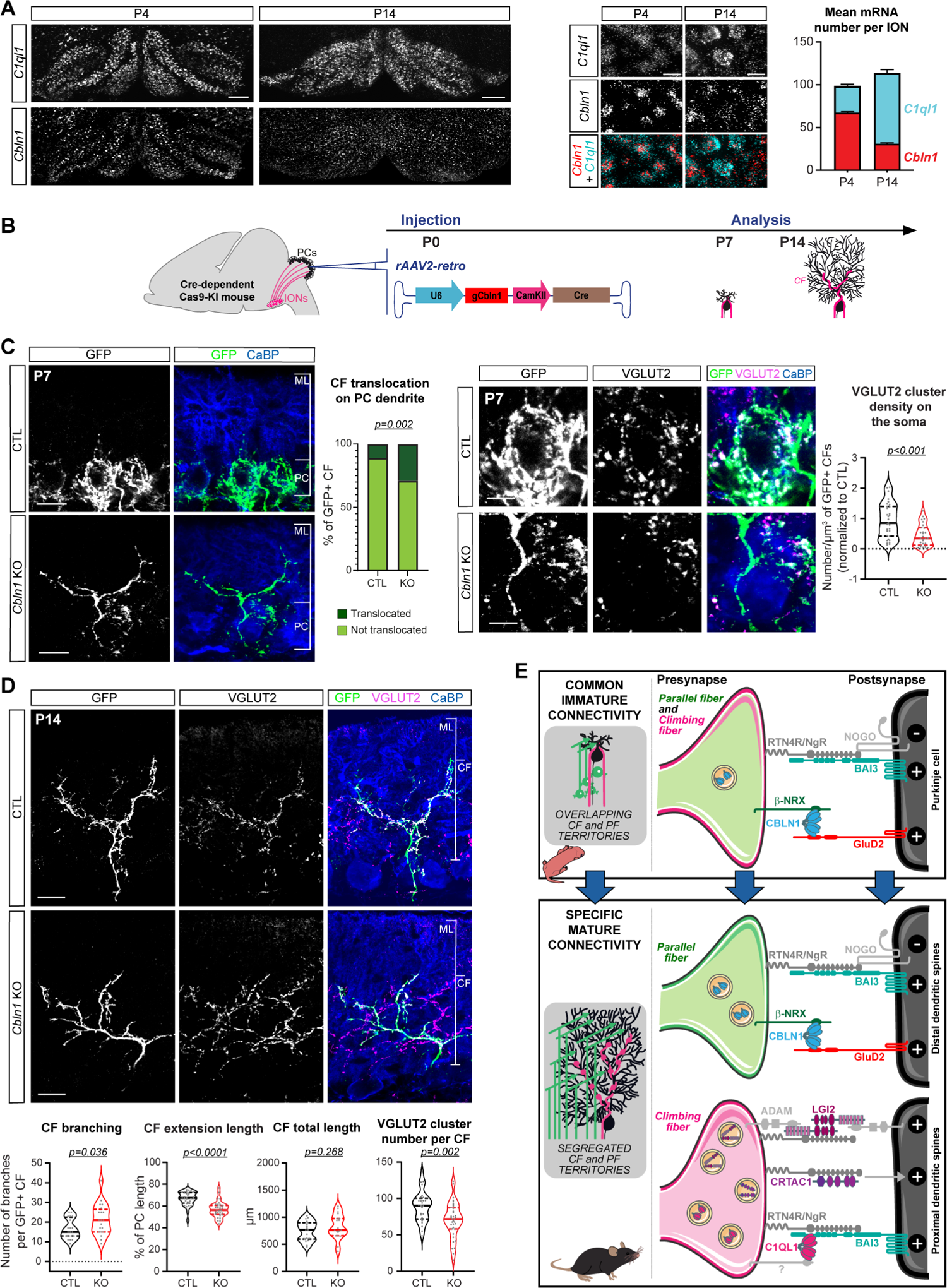
*Cbln1* expression in inferior olivary neurons is necessary for proper climbing fiber connectivity on Purkinje cells. **(A)** Duplex smFISH for *C1ql1* and *Cbln1* mRNAs in coronal sections from brainstem at P4 and P14. *Left panel:* Low magnification showing the inferior olive. *Right panel:* High magnification showing IONs and quantification of the mean number of both *C1ql1* and *Cbln1* mRNAs per ION at P4 and P14. Data are presented as mean ± SEM. n = 864-3910 cells, 4 animals per condition, 3 independent experiments. **(B)** Schematic illustration of the strategy of CRISPR/Cas9-mediated loss of function of *Cbln1* specifically in IONs. Stereotaxic injection of retrograde rAAVs expressing gRNAs targeting *Cbln1* or non-targeting control (CTL), under the U6 promoter, as well as the Cre recombinase, under the CamKII promoter, were performed in the cerebellum of Cre-dependent Cas9 knock-in (KI) mice at P0. Analyses of the morphology and connectivity of GFP positive (GFP+) CFs were performed at P7 or P14 in parasagittal cerebellar sections from CTL or *Cbln1* KO injected animals. **(C and D)** GFP+ CF and CF presynaptic boutons were immunostained for GFP (green) and VGLUT2 (magenta), respectively, and immunostaining for calbindin was used to label PCs and their dendritic tree (blue) in CTL and *Cbln1* KO. *(C) Left panel:* Quantification of the percentage of GFP+ CF translocated on PC dendrite in the molecular layer (ML). n≥ 87 CFs per condition. Statistics: Chi-square. Molecular layer (ML) extension and PC somata are indicated on the merge image. Scale bars, 20 µm. *Right panel:* High magnification of the GFP+ CF on the PC soma and quantification of the density of VGLUT2 clusters per GFP volume. Data are presented as violin plot with the median and quartiles. n ≥ 28 images, 8 animals per condition, 4 independent experiments. Statistics: unpaired Student’s test. Scale bars, 7 µm. *(D)* GFP+ CF morphology was assessed measuring 3 parameters: the number of branches per CF, the percentage of CF extension length on PC dendrite and the total length of each CF. VGLUT2 cluster number per GFP+ CFs was also quantified. Data are presented as violin plot with the median and quartiles. n = 20 CFs per condition, 6-8 animals per condition, 3 independent experiments. Statistics: unpaired Student’s test, Mann Whitney for CF branching. Scale bars, 20 µm. **(E)** Model of the change in the acquisition of molecular synapse identity of PC excitatory synapses during postnatal development. See also Figure S3.

## Discussion

Synaptome mapping using two excitatory postsynaptic markers has revealed that a large part of synapse diversity across brain regions is acquired during the first three postnatal months in rodents^3,36^. In particular, the first month is the stage of large increases in both numbers and types of synapses in many brain regions^36^. Analyzing the mechanisms regulating the development of excitatory connectivity on cerebellar PCs, we show that the increase in synapse diversity relies on specific molecular rules that control the formation of each synapse type on a given target neuron. GCs acquire their mature surfaceome, including the expression of synaptogenic genes, in one step during their differentiation, while IONs go through a multi-step definition of both their surfaceome and their mature connectivity on the PC target.

Many protein-receptor complexes have been shown individually to control synapse formation and maturation across the brain, including neurexins and their various ligands (LRRTMs, neuroligins…), secreted cerebellins and their GluD receptors, and secreted C1QL proteins and their receptors, the adhesion GPCRs of the BAI family^6,37^. However, to what extent does a combination of different signaling pathways contribute to defining the “cytochemical affinities” that underlie synapse specificity, as postulated by Sperry? Are these pathways functionally redundant for a single-synapse type? Our results demonstrate that at least five different molecular complexes contribute to CF/PC specific connectivity and identity (Figure 4E). Some molecular pathways regulate the development of mature connectivity at different levels: in axon and dendrite morphogenesis on one hand, and synapse formation and stabilization on the other hand. In the case of CFs, NRCAM and CBLN1 regulate in opposite direction CF morphogenesis in addition to regulating the number of synapses, in accordance with previous data^32,38,39^. The BAI synaptic receptors or the NOGO signaling pathway also control both morphogenesis and synaptogenesis in cerebellar PCs and other neuronal types across the brain^9,30,40,41^. Interestingly, we show that, in addition to the BAI3-ligand C1QL1, the cell surface proteins LGI2 and CRTAC1 control synapse numbers and maturation specifically without affecting CF morphogenesis. LGI2 is part of a family of secreted proteins that binds to ADAM receptors. CRTAC1 is an extracellular matrix protein that inhibits NOGO signaling and promotes synaptogenesis in cultured hippocampal neurons^42^. Thus, secreted proteins could play dual roles during synapse formation and specification: they could directly promote synaptogenesis *via* their specific receptors, but also inhibit other functions of their receptors by competing with other ligands, for example those regulating neuronal morphogenesis. Are these pathways redundant? Binding of LGI1 to NOGO receptors inhibits NOGO signaling and promotes binding of LGI1 to its ADAM receptor^43^. Recently interactions between the NOGO receptors and the BAI adhesion GPCRs were revealed to promote synaptogenesis in cultured human neurons^30^. This raises the possibility that C1QL1, like LGI and CRTAC1, also modulates NOGO signaling. These cell surface proteins could play partially redundant function by inhibiting NOGO signaling at the same synapse but in addition promote other signaling pathways *via* their specific receptors: BAI3 for C1QL1, ADAMs for LGI2 and other yet to be identified receptors for CRTAC1 (Figure 4E). This partial redundancy would be essential to maintain a certain robustness of circuit formation in the face of mutations in genes underlying synapse specificity.

Unexpectedly, both PC inputs rely initially on the same signaling pathway, *via* CBLN1 expression, for proper connectivity on their common target. The CBLNs constitute a subfamily of the C1Q family of secreted proteins that act as synaptic organizers throughout the brain^13,44,45^. CBLN1 is well known for promoting the formation and stabilization of PF/PCs synapses through the formation of a tripartite complex with presynaptic neurexin-1 and postsynaptic GluD2^11,20,46^. In addition, we show that CBLN1 promotes CF synapses on PC somata and that in its absence, the winning CF translocates faster on the growing PC dendritic arbor, but the number of synapses that it makes on the dendrites is still reduced. This correlates with previous observations that the receptor for CBLN1, GluD2, is transiently present at CF/PC synapses during development and adult reinnervation^47,48^. An original model postulated that the innervation of PCs on dendritic spines by PFs was a “default mode”^19^. Here, we show that a first step for PC connectivity is the recognition by the excitatory inputs, regardless of their type, *via* CBLN1/GluD2 complexes, constituting a sort of “default molecular code” for connectivity. Other genes that we found in our study could be part of this default molecular code such as the NOGO receptors that are found both in granule cells and immature IONs (supplemental data 4). During a second phase, molecular specification occurs selectively in the ION input: IONs start adding the expression of a specific combination of cell surface proteins that promotes the formation of synapses and their mature territory of innervation on the growing PC dendrites, in the face of competition with the PFs for their territory. Interestingly, many of the molecular pathways identified in our study are expressed in many regions of the brain and could control synapse specificity in other neuron types with synaptic organizations resembling the one observed for cerebellar PCs. Indeed, in the hippocampus, CA3 pyramidal neurons receive several types of excitatory inputs including one from dentate gyrus GCs that form a very specialized connection on the proximal dendrites *via* mossy fiber boutons. NRCAM, CRTAC1, LGI1 and C1QL proteins have been found at this particular synapse^49,50^. It is thus tempting to postulate that similar molecular rules control the development of excitatory connectivity in other neurons across the brain. For example, is the molecular identity of dentate gyrus-CA3 synapses acquired actively during development, while other synapses made on CA3 hippocampal neurons rely on a “default molecular code”?

What regulates the multi-step acquisition of the molecular identity of certain synapse types while in others synapse molecular specification occurs in a single step? The transcriptional program underlying the change in the gene expression pattern would need to be controlled by either an internal program or actively regulated. If it is active, it could be due to molecular factors, as recently described in Drosophila^51^, or due to changes in neuronal activity in the network^52^. The postnatal developmental period of circuit formation would thus be very sensitive to modulation by external factors for certain types of synapses but not others. This would imply specific consequences on circuit development by environmental changes in terms of computation and symptom etiology. The facts that many human brain disease genes code for synaptic proteins, and that synapse diversity can be modulated by mutations relevant to neurodevelopmental disorders, highlight the importance of understanding the molecular rules generating synapse diversity during development^53,54^.

## EXPERIMENTAL PROCEDURES

### Animals and AAV injections

All animal protocols and animal facilities were approved by the Comité Régional d’Ethique en Expérimentation Animale (#22406) and the veterinary services (D-75-05-12). OF1 (Charles River Laboratories, Wilmington, USA) were used for the determination of developmental expression patterns or smFISH. The Cre-dependent Cas9-KI mouse line (Cas9-KI) was obtained from The Jackson Laboratory (B6J.129(B6N)-Gt (ROSA)26Sor^tm1(CAG-cas9*,-EGFP)Fezh^/J, strain #026175). The NeuroD1Cre mouse line (Tg(Neurod1-cre)RZ24Gsat/Mmucd) was obtained from Pr. Mary Beth Hatten (The Rockefeller University, USA). Both lines are maintained on the C57Bl6/J background, and homozygous mice were crossed with C57Bl6/J wild-type animals to generate heterozygous animals used in our experiments.

Injections of AAV particles were performed in the cerebellar vermis of ice-anesthetized P0 Cas9-KI heterozygous mice at 1 mm depth from the skull and at 3.2 mm relative to Bregma, to target Purkinje cell layer. AAV2/retrograde serotype was used to target IONs. 0.25µl of AAV was injected per animal using pulled calibrated pipets.

### Gene expression analysis

#### High-throughput RTqPCR and ddPCR™

were performed as previously described ^23^ (see also supplemental methods). Unsupervised clustering of log2 relative gene expression during postnatal development was performed using Cluster 3.0 program. Each gene expression value was adjusted, so that the median (more robust against outliers) value of each raw is 0, to reflect their variation compare to the *Rpl13a* reference gene. Agglomerative hierarchical clustering was performed using complete linkage to compute the distance matrix between gene expression data. Hierarchical clustering methods assemble genes and time points in a tree structure, where the size of the branches increases when their similarity decreases to each other. Treeview allowed the visualization of the data.

#### Single molecule Fluorescent *In situ* Hybridization (smFISH)

Animals at P4, P14 or P>60 (adult) were processed for smFISH labeling using the RNAscope Multiplex Fluorescent Assay kit (Advanced Cell Diagnostics, Newark, USA) according to manufacturer’s instructions with minor modifications as detailed in the supplemental methods. Probes were designed to target all predicted transcripts variants for each candidate gene.

### Immunohistochemistry

Immunostaining were performed on 30 µm-thick parasagittal cerebellar sections obtained using a freezing microtome after intracardiac perfusion of mice with fresh 4% paraformaldehyde in phosphate buffer saline (PBS). Sections were blocked with 4% normal donkey serum in PBS containing 1% Triton X-100 for 45 min at room temperature. The primary antibodies (see supplemental material) were diluted in PBS supplemented with 1% Triton X-100, 1% donkey serum, and incubated two hours at room temperature.

### Image acquisition

All image stacks were acquired using a spinning-disk confocal CSU-W1 microscope with a 25X or 63X objective. The mosaic images of smFISH experiments were reconstructed using the Metamorph software (Figure 1E and 4A, left panel). Quantitative analysis of images was performed using custom-made plugins as described in the supplemental methods.

### Statistical analysis

All statistical analyses were performed using GraphPad Prism8. Normality was assessed using D’Agostino and Pearson, Shapiro-Wilk or Kolmogorov-Smirnov normality tests. When groups fit to the normal distribution, the differences between the two groups were assessed using two-tailed Student’s t-test. When the populations did not fit the normal distribution, the non-parametric Mann-Whitney test was used. Differences in distribution were tested using the chi-squared test. Data are presented as mean ± SEM, or as violin plots with median and quartiles.

---

**Figure S1.**
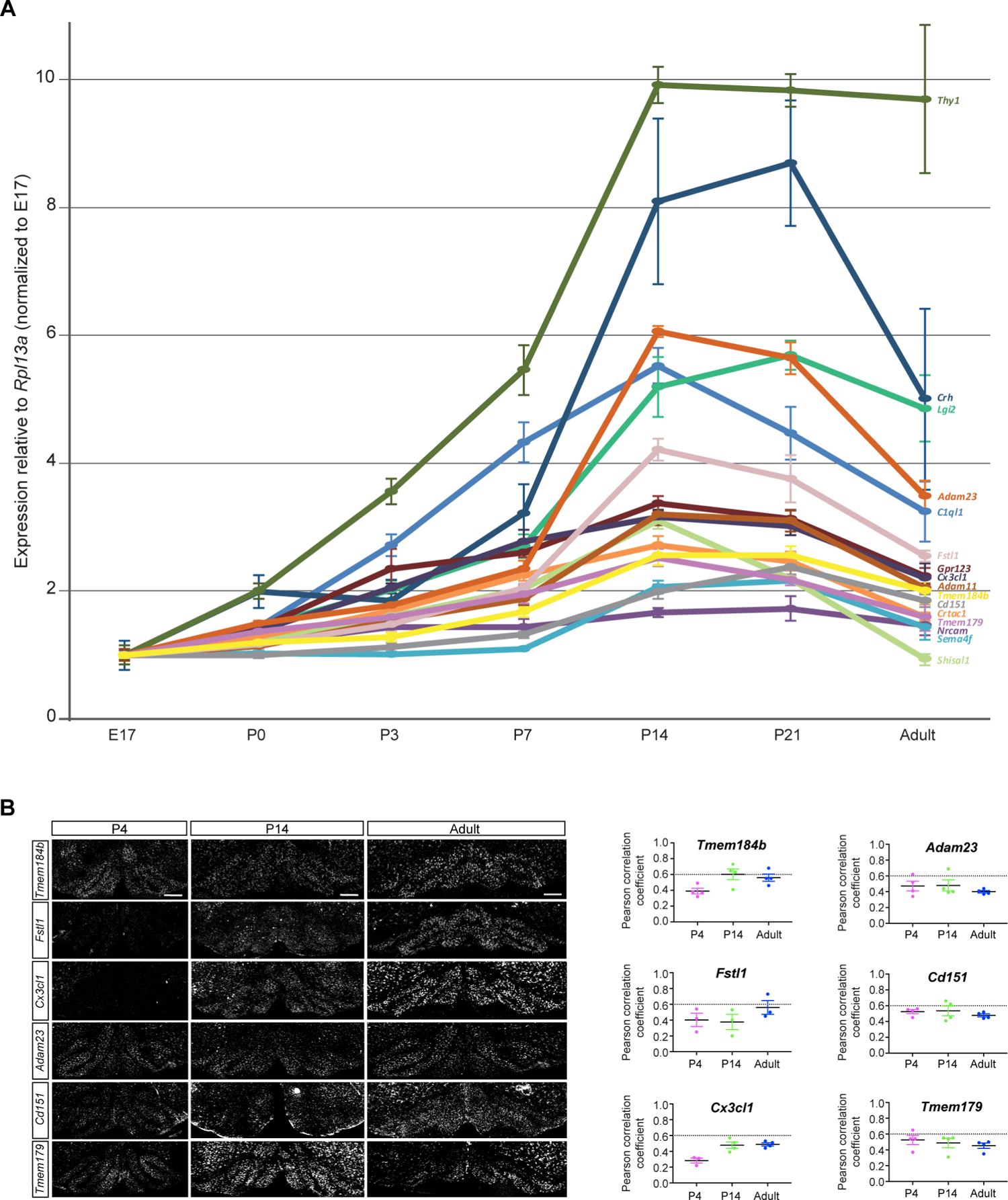
Developmental expression pattern of ION candidate genes. **(A)** Expression pattern of the fifteen ION DEGs selected as candidate and assessed using high-throughput RT-qPCR on brainstem RNA extracts at different stages of development (Embryonic day, E17, Postnatal days, P0, P3, P7, P14, P21) and in adult. Expression levels were normalized to *Rpl13a* gene. Data are presented as mean ± SEM. n = 4 animals per stage. **(B)** *Left panel:* duplex smFISH for *C1ql1* and six candidate mRNAs in coronal sections of the brainstem at P4, P14 and in adult. *Right panel:* correlation between *C1ql1* and candidate mRNAs were determined at the brainstem level using the Pearson correlation coefficient, revealing ION DEGs slightly correlated with *C1ql1* (coefficient >0.6 corresponding to high correlation). Data are presented as mean ± SEM. n = 3-4 animals per condition, 2-3 independent experiments. Scale bars, 150 µm.

**Figure S2.**
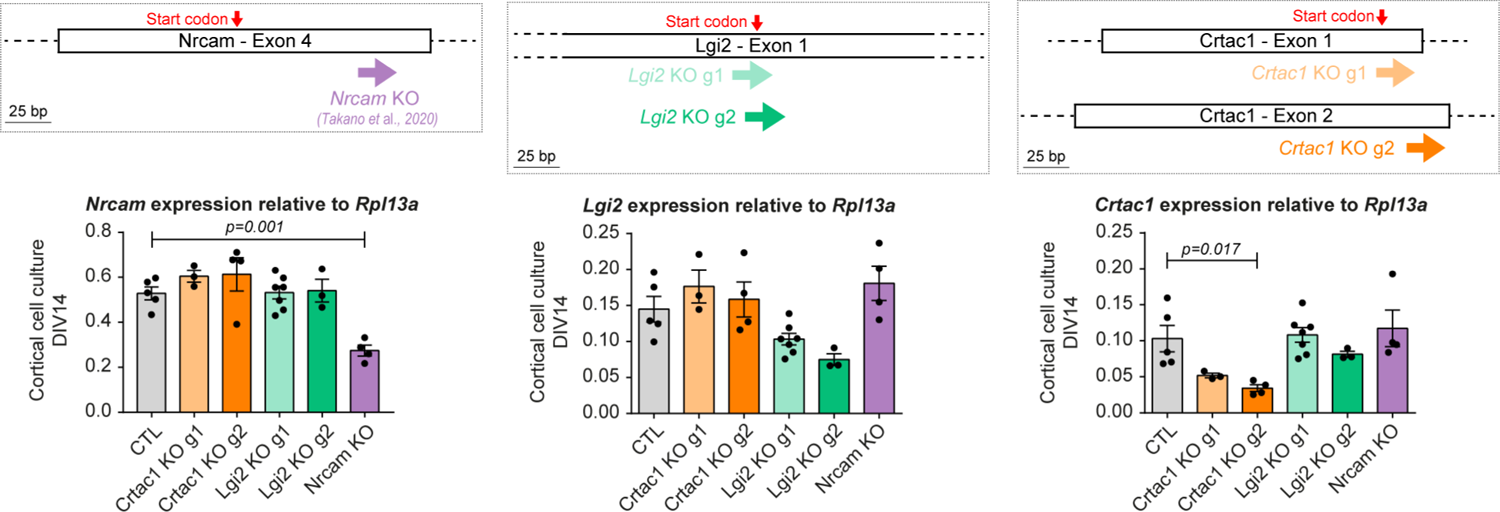
Validation of CRISPR/Cas9 gRNAs for *Nrcam, Lgi2* and *Crtac1* KO. *Top panel:* illustration of the genomic sequence of mouse *Nrcam, Lgi2* and *Crtac1* with the location of the CRISPR/Cas9 gRNAs tested for each gene. *Bottom panel:* To test CRISPR/Cas9 efficiency, expression of *Nrcam, Lgi2* and *Crtac1* mRNAs was assessed using quantitative RT-PCR on RNA extracts from cortical cell cultures transduced at 3 days *in vitro* (DIV3) with AAVs driving the expression of each gRNA directed against candidate genes or non-targeting control gRNA (CTL). Analysis was done at DIV14. Expression levels were normalized to the *Rpl13a* gene. Data are presented as mean ± SEM. n = 3-7 independent experiments. Statistics: one-way ANOVA with multiple comparisons.

**Figure S3.**
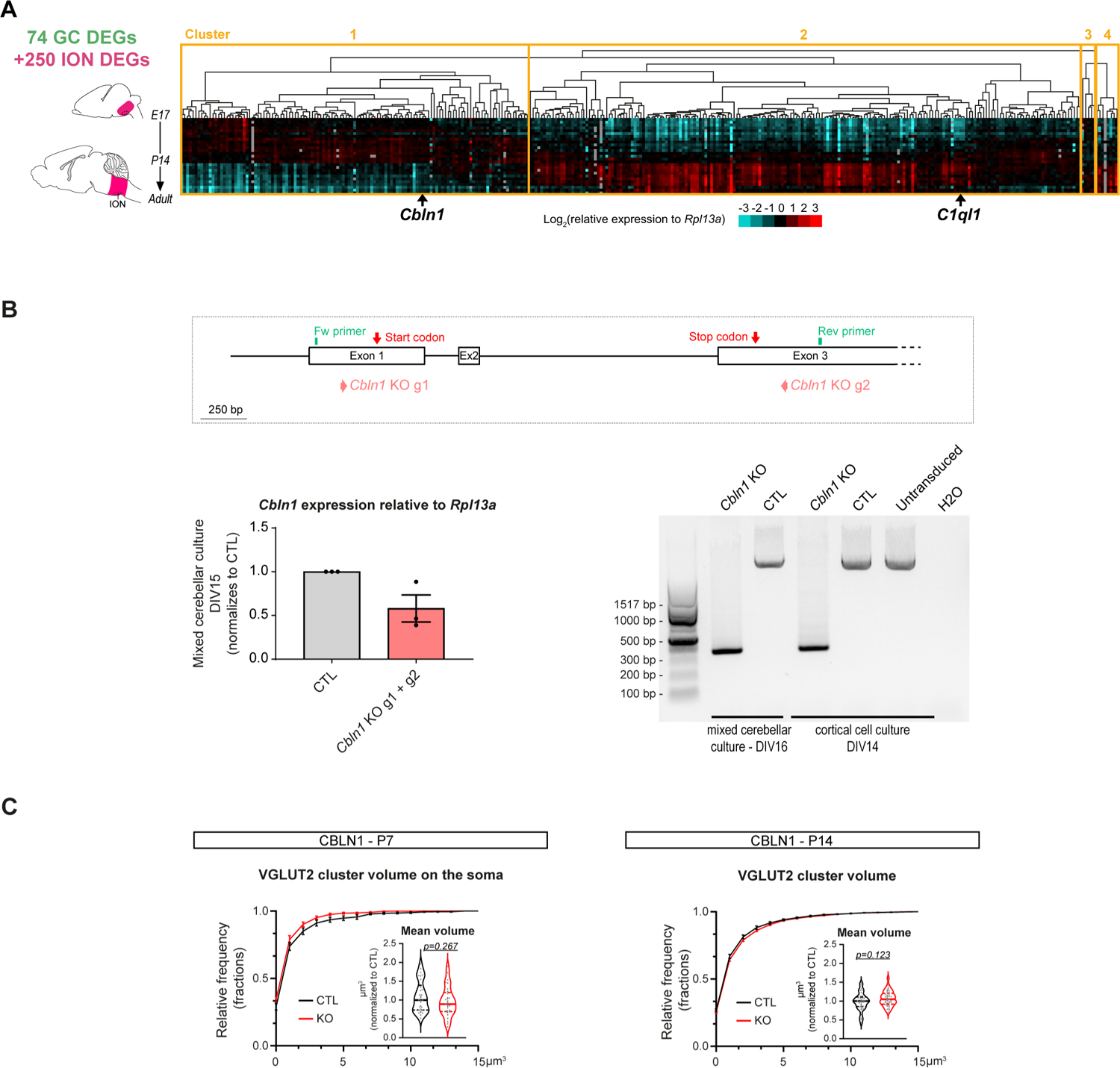
Cbln1 expression dynamics in the brainstem and validation of CRISPR/Cas9 gRNAs for *Cbln1* KO. **(A)** Heatmap of the 74 GC DEGs and 250 ION DEGs in the brainstem between E17 and adult revealing *Cbln1* as an early-expressed gene. **(B)** *Top panel:* illustration of the genomic sequence of mouse *Cbln1* with the location of the CRISPR/Cas9 KO guide RNAs (*Cbln1* KO g1 and g2) as well as forward (Fw) and reverse (Rev) primers targeting non-coding regions of *Cbln1* exon1 and exon 3, respectively. *Bottom left panel:* to test CRISPR/Cas9 efficiency, the expression of *Cbln1* mRNAs was assessed using quantitative RT-PCR on RNA extracts from mixed cerebellar cultures transduced at days *in vitro* 2 (DIV2) with AAVs driving the expression of *Cbln1* gRNAs or non-targeting control gRNA (CTL) Analysis was done at DIV15. Expression levels were normalized to *Rpl13a* gene. Data are presented as mean ± SEM. n = 3 independent experiments. *Bottom right panel:* CRISPR/Cas9 efficiency was tested by PCR analysis of endogenous *Cbln1* in purified genomic DNA from mixed cerebellar cultures or cortical cell cultures at DIV16 or DIV14, respectively. Transduction was performed at 2 days *in vitro* (DIV2) for cerebellar mixed cultures and DIV3 for cortical cell cultures with AAVs driving the expression of each gRNA directed against *Cbln1* or non-targeting control gRNA (CTL). Wild-type size of the fragment is expected at 2663 bp and KO fragment at 380 bp. **(C)** Distribution and mean of the VGLUT2 cluster volume, presented as violin plot, were quantified in GFP+ CFs both at P7 and P14 in CTL and *Cbln1* KO. P7 VGLUT2 cluster volume: n ≥ 29 images per condition; 8 animals per condition, 4 independent experiments; statistics: Mann-Whitney. P14 VGLUT2 cluster volume: n ≥ 33 images per condition; 8-9 animals per condition, 3 independent experiments; statistics: unpaired Student’s test.

## Acknowledgments

We would like to thank Adeline Boyreau for her help with some of the experimental procedures, France Maloumian for her help with infographics and the personnel from the CIRB animal and imaging facilities. In particular, we thank Héloïse Monet for her help in developing the plugin for VGLUT2 quantification. High throughput qPCR was carried out on the qPCR-HD-Genomic Paris Centre platform supported by grants from Région Ile-de-France. This work was supported by funding from: Fondation pour la Recherche Médicale Equipe FRM DEQ20150331748 (FS), European Research Council ERC consolidator grant SynID 724601 (to FS), Q-life ANR-17-CONV-0005 (to FS), ANR-10-LABX-54 MEMO LIFE (to FS), Sorbonne Université ED158 (to MP), Collège de France (to MP).

## Notes

### Competing Interest Statement

The authors have declared no competing interest.

### Summary of Updates

This version of the manuscript has been revised to update all figures and supplemental figures that were on a transparent background and are now on a white background.

